# A Commercial ARHGEF17/TEM4 antibody cross-reacts with Nuclear Mitotic Apparatus protein 1 (NuMA)

**DOI:** 10.1101/2022.05.10.491389

**Authors:** Diogjena Katerina Prifti, Annie Lauzier, Sabine Elowe

## Abstract

The Rho family Guanine nucleotide exchange factor (GEF) ARHGEF17 (also known as TEM4) is a large protein with only 3 annotated regions: an N-terminal actin-binding domain, a Rho-specific dbl homology (DH)- pleckstrin homology (PH) type GEF domain and a seven bladed β propeller fold at the C-terminus with unknown function. TEM4 has been implicated in numerous activities that rely on regulation of the cytoskeleton including cell migration, cell-cell junction formation and the spindle assembly checkpoint during mitosis. Here we have assessed the specificity of a TEM4 polyclonal antibody that has been commonly used as a Western blotting and immunocytochemistry probe for TEM4 in mammalian cells. We find that this antibody, in addition to its intended target, cross-reacts with the Nuclear Mitotic Apparatus Protein 1 (NuMA) in Western blotting and immunoprecipitation, and detects NuMA preferentially in immunocytochemistry. This cross-reactivity, with an abundant chromatin- and mitotic spindle-associated factor, is likely to affect the interpretation of experiments that make use of this antibody probe, in particular by immunocytochemistry and immunoprecipitation.

## Introduction

Signaling events mediated by Rho family GTPases orchestrate cytoskeletal dynamics that direct many aspects of the cell including its morphology, motility, adhesion, endocytosis, cell cycle progression and cytokinesis, apoptosis and survival (1, 2). The Rho family guanine nucleotide exchange factor ARHGEF17 (also known and subsequently referred to as Tumour Endothelial marker 4, TEM4) plays an important role in the regulation of the interphase cytoskeleton. In endothelial cells, TEM4 promotes the persistence of cellular migration by regulating actin stress fibre structures and cellular adhesions (3). Indeed TEM4 can directly associate with actin through an actin binding sequence found in its N-terminus, and independently of its RhoGEF activity (3). In agreement, full-length TEM4 or constructs including the N-terminal actin binding domain expressed in Human umbilical vein endothelial cells (HUVECs) or NIH/3T3 (isolated fibroblast from mouse NIH/Swiss embryo) cells localized strongly to the actin structures including stress fibres (3). In vitro, the actin binding domain showed a strong preference for dynamic, newly assembled f-actin filaments. In concordance with a function in regulating cytoskeletal dynamics, TEM4 is also implicated in the maintenance of cell-cell adhesion and junctions (4) and depletion of endogenous TEM4 by shRNAs impaired Madin–Darby canine kidney (MDCK) and HUVEC cell junctions, disrupted MDCK actin formation in 3D culture and resulted in disfunction of endothelial cells barriers (4). In an immunocompetent mouse model, TEM4 was found to be involved in tumor growth and metastatic dissemination of lung cancer cells (5). Most recently, the Mitocheck genome wide RNAi screening effort had identified TEM4 as an essential mitotic protein (6). Subsequent studies proposed that TEM4 may function as a new player in the spindle assembly checkpoint during mitosis functioning as a timer for retention of the critical checkpoint kinase Monopolar Spindle 1 (MPS1) (7). Importantly, these studies collectively based their conclusions on both manipulation of the endogenous protein as well as overexpression of various TEM4 fragments. Overall, these data support the idea that TEM4 is an important regulator of cellular cytoskeletal structures and cell cycle progression.

Antibodies are critical tools for biomedical research. However, their value depends on faithful characterization of both affinity and specificity for the intended target. The validation of antibodies has unfortunately significantly faltered compared to their production, resulting in only very basic and non-quantitative verification, followed by the consequent flaws in the reliability of many of the available reagents (Baker,2020, 2015). One important issue is cross-reactivity due to off- target binding (10–15). In particular, the ratio of the target protein to other proteins in a sample may lead to significantly different levels of off-target binding depending on the application. This consideration may be particularly relevant when dealing with concentrated pools of proteins at distinct subcellular locations, even if the antibody’s affinity for such off-targets is considerably lower than its affinity for the target protein. Another issue is that sample preparation is substantially different depending on the intended application, which can influence the epitopes exposed on the target protein. Thus, antibodies must be validated in an application-specific manner, as was recently suggested by the International Working Group for Antibody Validation (IWGAV) (16).

Here, we evaluate the specificity of a commercial rabbit polyclonal anti-TEM4 antibody available from multiple providers and that is used in the literature in multiple applications including Western blotting and immunocytochemistry (3–5,7,17). Our results reveal strong cross-reactivity of this antibody with the Nuclear Mitotic Apparatus Protein 1 (NuMA) in Western blotting, immunoprecipitation and immunocytochemistry, indicating major limitations of its utility in these approaches.

## RESULTS

### TEM4 antibody staining shows spindle localization

We initially sought to investigate the function of TEM4 in mitosis. TEM4 had been previously shown to be an essential mitotic protein that localizes to kinetochores that is likely involved in generating a robust spindle assembly checkpoint (6, 7). We first performed immunofluorescence using a commercially available polyclonal antibody commonly used in the literature to study this protein (RRID: AB_1141641). Surprisingly, in HCT-116 cells in metaphase, we observed strong mitotic spindle staining that did not appear to significantly overlap with kinetochores, as indicated by poor co-localization with the CREST anti-centromere marker (Fig. 1A). To further explore immunofluorescence signals detected by the TEM4 antibody, we stained HCT-116 cells in interphase and across the multiple stages of mitosis. In interphase cells, we observed distinct nuclear staining whereas from prometaphase to anaphase TEM4 antibody signals strongly resembled spindle staining. In telophase, the TEM4 antibody decorated the segregating chromatin masses, suggesting reincorporation of the epitope into the nucleus (Fig. 1B, supp Fig.1). To rule out the idea that spindle localization could be cell-line specific, we stained a panel of human immortalized and cancer cells lines commonly used for the study of mitosis (HCT-116, HeLa, HeLa S3, RPE1-hTERT and U2OS cells) under the same conditions. All five cell lines showed clear spindle staining in mitotic cells using the TEM4 antibody (Fig. 1C).

**Fig. 1.**
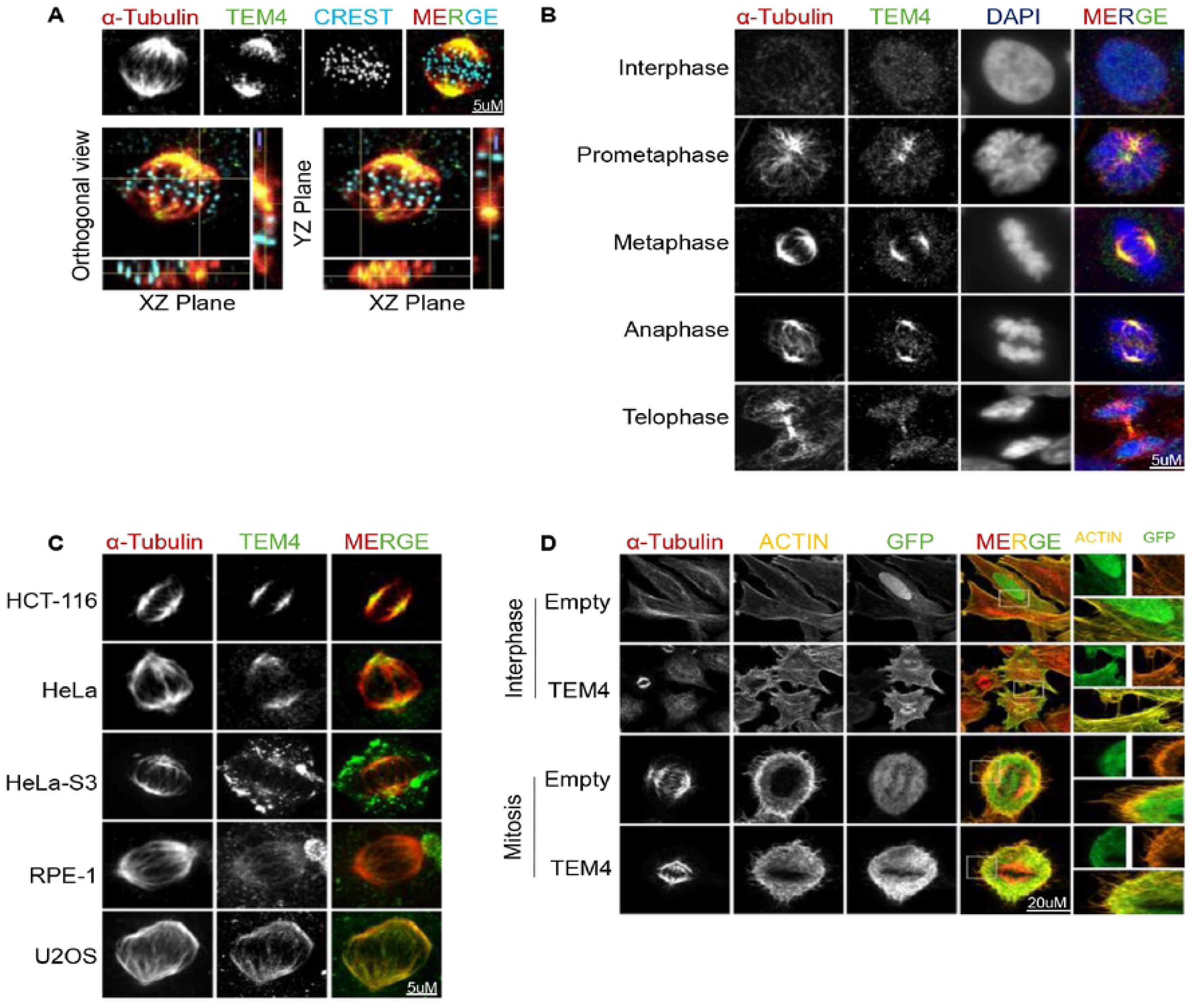
The TEM4 antibody decorates the mitotic spindle. A: Representative immunofluorescence image of a metaphase HCT-116 cell. Cells were synchronized in mitosis after 10 hours of release from double thymidine block and fixed for 10 minutes with PTEMF buffer. Fixed cells were stained with anti-TEM4 (green), anti-α-tubulin for the spindle microtubules (red) and CREST serum for the kinetochores (blue). In the lower panel, orthogonal sections of the maximum intensity projections of the same cell are shown. Scale bar = 5μm. B: Representative immunofluorescent images of TEM4 antibody staining in HCT-116 cells during cell cycle progression. Fixed cells were stained with anti-TEM4 (green), anti-α-tubulin (red), and DNA was stained with Hoechst (blue). Cell cycle stages are indicated to the left of each image and merged images are shown in the right most panel. Scale bar = 5μm C: Immunofluorescence of metaphase spindles in different cell lines lines showing staining at the mitotic spindle. Cells were synchronized in mitosis after 10 hours of release from double thymidine block and fixed for 10 minutes with PTEMF buffer. Fixed cells were stained with anti-TEM4 (green) and anti-α-tubulin for the spindle microtubules (red). The different cell lines are indicated to the left of each image. Scale bar = 5μm D: HeLa T-REx cells were left synchronous (upper panel) or were synchronized in mitosis (lower panel) after 30 minutes release from RO-3306. Cells expressing empty-GFP, or GFP-TEM4 were stained with anti-GFP (green), anti-α-tubulin for the spindle microtubules (red) and phalloidin-Atto 565 for F-actin (Yellow). Scale bar = 20μm.

The observed nuclear staining in interphase was somewhat surprising, given that previous results demonstrated localization to the actin cytoskeleton and cell junctions of overexpressed, tagged full-length TEM4 in Human umbilical vein endothelial cells (HUVECs), HeLa and MDCK cell lines, (4, 17). In line with these observations, we also found that GFP-TEM4 localized to the actin cytoskeleton in interphase HeLa cells and to the actin-rich cortex in mitosis (Fig. 1D). Moreover, we observed no clear colocalization with microtubule structures. Importantly, changing the position of the GFP-tag to the c-terminus or indeed a using a different tag (we tested flag), gave similar results suggesting that the tag itself does not influence localization (data not shown). Overall, these observations indicate that while exogenously expressed TEM4 localized predominantly to actin structures during both interphase and mitosis, the TEM4 polyclonal antibody primarily stained the nucleus in interphase and the spindle during mitosis.

### A Peptide competition assay confirms reactivity of the TEM4 antibody to the mitotic spindle

Previous observations have highlighted the localization and function of several RhoGEFS at the spindle during mitosis suggesting that TEM4 may indeed be a spindle-associated protein (18–21). We therefore considered the possibility that the endogenous and ectopic versions of TEM4 may exhibit different localization and behavior. In order to test this idea, we first attempted to rule out non-specific spindle staining by the TEM4 antibody. To this end, we performed a competition assay where we blocked the TEM4 antibody with a peptide corresponding to the 18 amino acid epitope located within residues 910-960 of TEM4. Preincubation of the TEM4 antibody with the immunogenic peptide but not vehicle control resulted in dramatic loss of spindle staining in mitotic HCT-116 cells (Fig. 2A, B). Similarly, in Western blots of total cell lysates from HeLa S3 cells, preincubation of the TEM4 antibody with the blocking peptide led to disappearance a 220 kDa band corresponding to the predicted molecular weight of full-length TEM4 whereas this band was readily detected in the same lysates subjected to Western blotting with the TEM4 antibody in the absence of the blocking peptide (Fig. 2C). Of interest, preincubation with the antigen also inhibited antibody reactivity with a second prominent band of approximately 180kDa in both HeLa S3 and HCT-116 cells. Collectively, these data indicate that the commercial TEM4 antibody specifically reacts with a mitotic spindle associated epitope by immunofluorescence and with two major protein species at 220kDa (presumably TEM4) and 180 kDa by Western blotting.

**Fig. 2.**
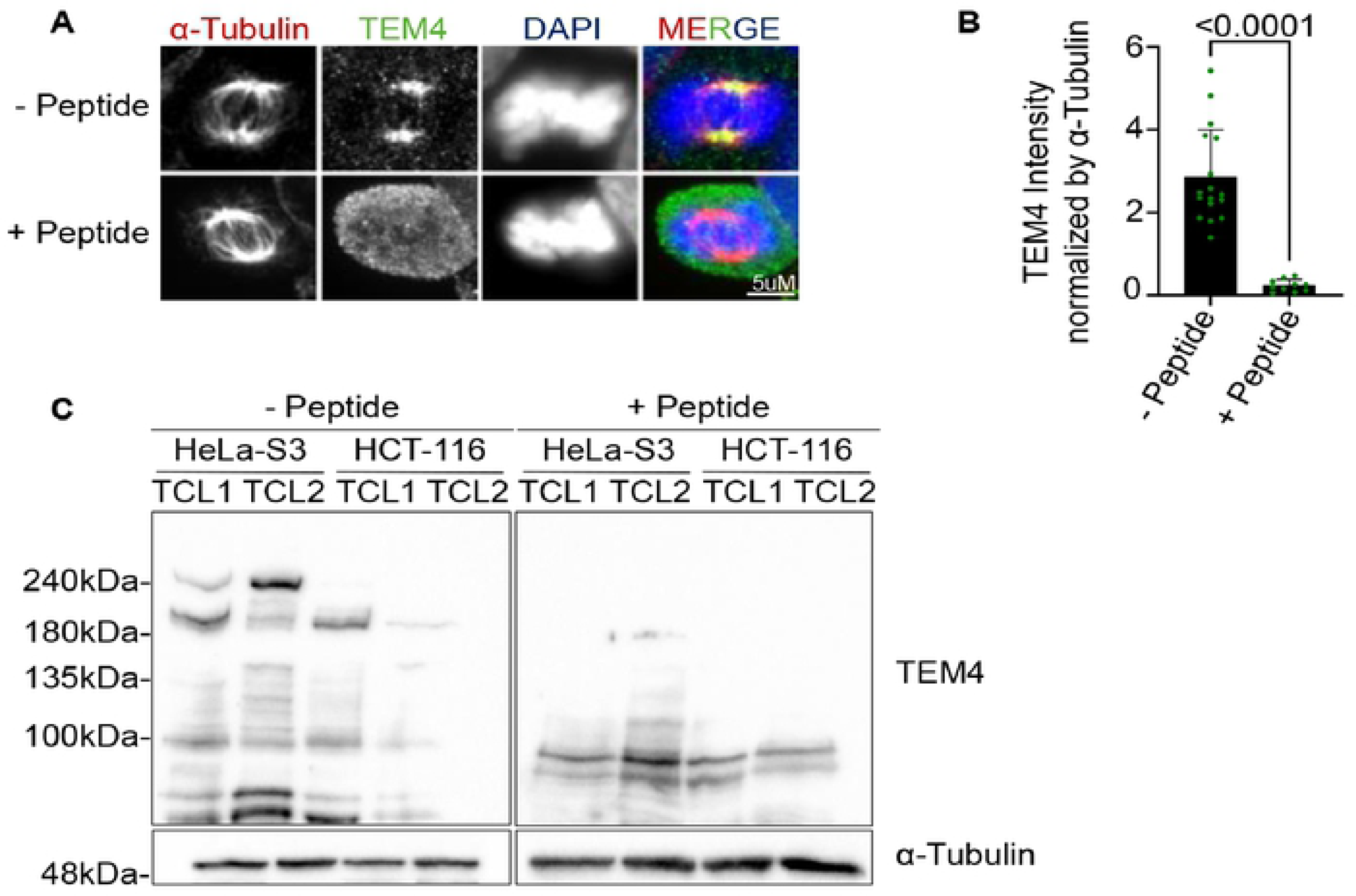
A Peptide competition assay confirms reactivity of the TEM4 antibody to the mitotic spindle. A: TEM4 blocking peptide competition assay in HCT-116 cells. Cells synchronized in mitosis after 10 hours of release from a double thymidine block and fixed for 10 minutes with PTEMF buffer. TEM4 antibody (1 μg) was pre-incubated with either vehicle control (top panel) or with the immunogenic peptide (20 μg) corresponding to the 18 amino acid epitopes within the 910-960 residues of TEM4 (bottom panel). Cells were stained with anti-TEM4 (green), anti-α-tubulin for the spindle microtubules (red) and and DNA is stained with HOECHST (blue). Scale bar = 5μm. B: Statistical analysis of TEM4 intensity at the mitotic spindle incubated either with vehicle control or with the immunogenic TEM4 peptide from A. The experiment was performed in one biological replicate and the P value is calculated based on the mean SD of cells per condition. C: Immunoblots of HeLa S3 or HCT-116 cell lines. Total cell lysates from both cell lines were extracted with RIPA buffer and run on a polyacrylamide gel. Western blot was performed with either TEM4 (1 μg) antibody incubated with vehicle control (left panel) or with the immunogenic TEM4 peptide (20 μg) (right panel).

### The TEM4 antibody targets microtubule minus ends

We were particularly intrigued by the potential localization of TEM4 to spindle microtubules. To investigate this in more detail, we asked whether TEM4 antibody staining changes with alterations to microtubule stability. Treatment of HCT-116 cells with the microtubule destabilizing reagent nocodazole resulted in loss of the spindle and concomitant loss of TEM4 antibody reactivity. On the other hand, hyperstabilized microtubules as a result of Taxol treatment resulted in multipolar spindles as shown before (22, 23) with a strong TEM4 antibody staining (Fig. S2A). Quantification of TEM4 antibody staining showed a relative increase in signal intensity at multipolar spindles in the presence of Taxol (Fig. S2B).

To identify more precisely the localization of the TEM4 antibody target within the mitotic spindle, we sought to identify the precise population of microtubules that was detected by this antibody. We first considered the possibility that the TEM4 antibody associated with stably attached kinetochore fibres (k-fibres). To test this, cells were allowed to enter mitosis in the presence of MG132 which allows for the congression and alignment of chromosomes at metaphase with stable attachments between kinetochores and k-fibres before incubation at 4°C for 10 minutes to depolymerize poorly-attached microtubules. In this assay, k-fibres forming stable end-on attachments to kinetochores are maintained whereas lateral attachments between kinetochores and microtubules, unattached microtubules and interpolar spindle microtubules are destabilized and depolymerized (24). K-fibers retained after cold treatment of HCT-116 cells exhibited a visible interaction to kinetochores (marked by CREST, Fig. 3A). In these cells, and in agreement with our observations in Fig. 1A, TEM4 antibody staining was not detected in the proximity of kinetochore, rather becoming focused towards the minus end of microtubules. Measurement of overall TEM4 antibody signal intensity indicated a strong reduction in signal in cells incubated at 4°C when compared to cells maintained at 37°C (Fig. 3B).

**Fig. 3.**
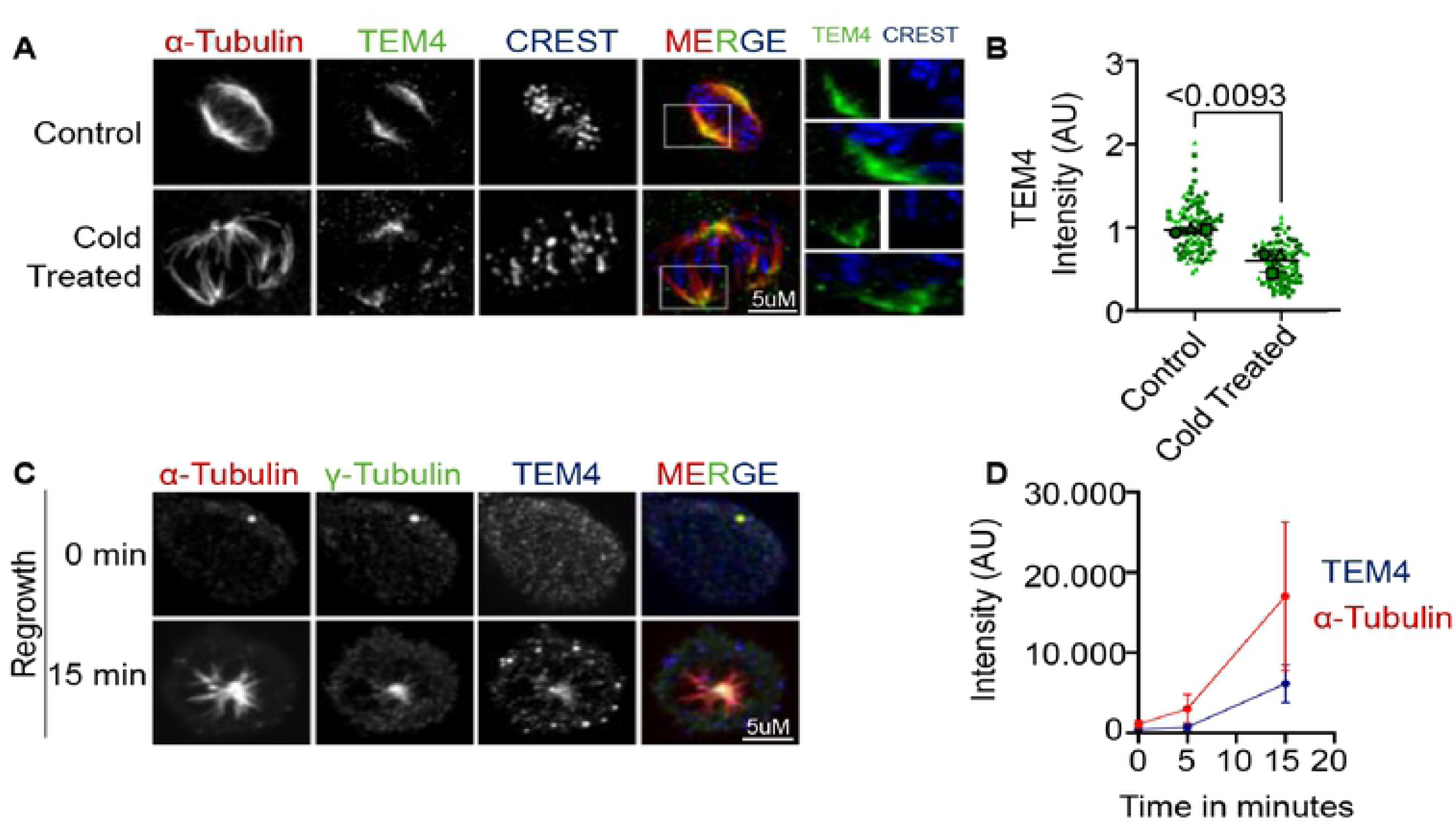
TEM4 antibody detects a target localized to microtubule minus ends. A: Representative Immunofluorescent image of mitotic spindles depolymerized by cold treatment. Control (upper panel) or cold treated for 20 minutes HCT-116 cells (bottom panel) fixed with PTEMF solution and stained with anti-TEM4 (green), anti-α-tubulin for the spindle microtubules (red) and CREST serum for the kinetochores (blue). Scale bar = 5μm. B: Statistical analysis of TEM4 intensity at the mitotic spindle in cells from A. A minimum of 20 cells was analyzed per condition for a total of 3 independent experiments. The graph shows superplots where each biological replicate is color coded and the averages from each replicate are indicated accordingly. For all protein intensity measurements at the mitotic spindle, each dot represents a cell, and the error bars display the variation between the mean of biological replicates. Two-tailed, non-paramentrric Mann-Whitney test t-test was performed to compare the mean values between the tested groups. C: Immunofluorescent image HCT-116 cells treated for three hours with 3μM of Nocodazole. Cells were fixed with methanol after Nocodazole was washed out and cells were incubated with fresh media for the indicated time points. Fixed cells were stained with anti-TEM4 (blue), anti-α-tubulin for the spindle microtubules (red) and anti-γ-tubulin for the centrosomes (green). Scale bar = 5μm. D: Statistical analysis of α-tubulin and TEM4 intensity in different time points corresponding to figure C.

The observation that TEM4 antibody decorated microtubules distant from kinetochores suggests that it detects an epitope at microtubule minus ends. To test this idea, we stained growing microtubules with the TEM4 antibody. As expected, when spindle regrowth was observed after depolymerization by Nocodazole, microtubules (marked by α-tubulin) repolymerized in a radial manner, forming a distinct aster-like structure. TEM4 antibody signal overlapped with and tracked γ-tubulin, a minus-end marker and an essential component of centrosomes and the γ-tubulin ring complex (25–31) (Fig. 3C, D). Collectively, these data strongly suggest that the TEM4 antibody detects a target localized to microtubule minus ends.

### TEM4 and NuMA localize to similar microtubule structures

The dominant microtubule nucleation pathway for spindle formation in mitosis is through a centrosome mediated pathway in cells where these organelles exist, although it is widely accepted that acentrosomal microtubule assembly in the cytoplasm and at kinetochores both contribute significantly to efficient chromosome attachment and congression (32–36). To determine whether the TEM4 antibody associated with both centrosomal and non-centrosomal microtubules, HeLa cells were treated with the 100 nM PLK4 (Polo-like kinase 4) inhibitor, centrinone, to specifically inhibit centrosomes duplication and generate acentrosomal cells (37). As previously shown, loss of centrosomes resulted in multiple γ-tubulin clusters including bundle-like and monopolar- like clusters compared to the defined centrosomes seen in control cells (Fig. 4A) (37). In addition to staining near the spindle poles, TEM4 antibody staining revealed a bundled structure between the poles with striking similarity to the recently reported structures formed by the microtubule minus end focusing protein, NuMA (38) (Fig. 4A). The same structures were observed after a 10 minute cold treatment (Fig. S2C, D). To confirm that the observed structures coincided with NuMA, we co-stained bipolar control cells and acentrosomal cells generated after centrinone treatment with TEM4 and NuMA antibodies which demonstrated appreciable colocalization of the signals in these interpolar bundles (Fig. 4B). Overall, our observations here support the localization of TEM4 to NuMA-rich interpolar bundles.

**Fig. 4.**
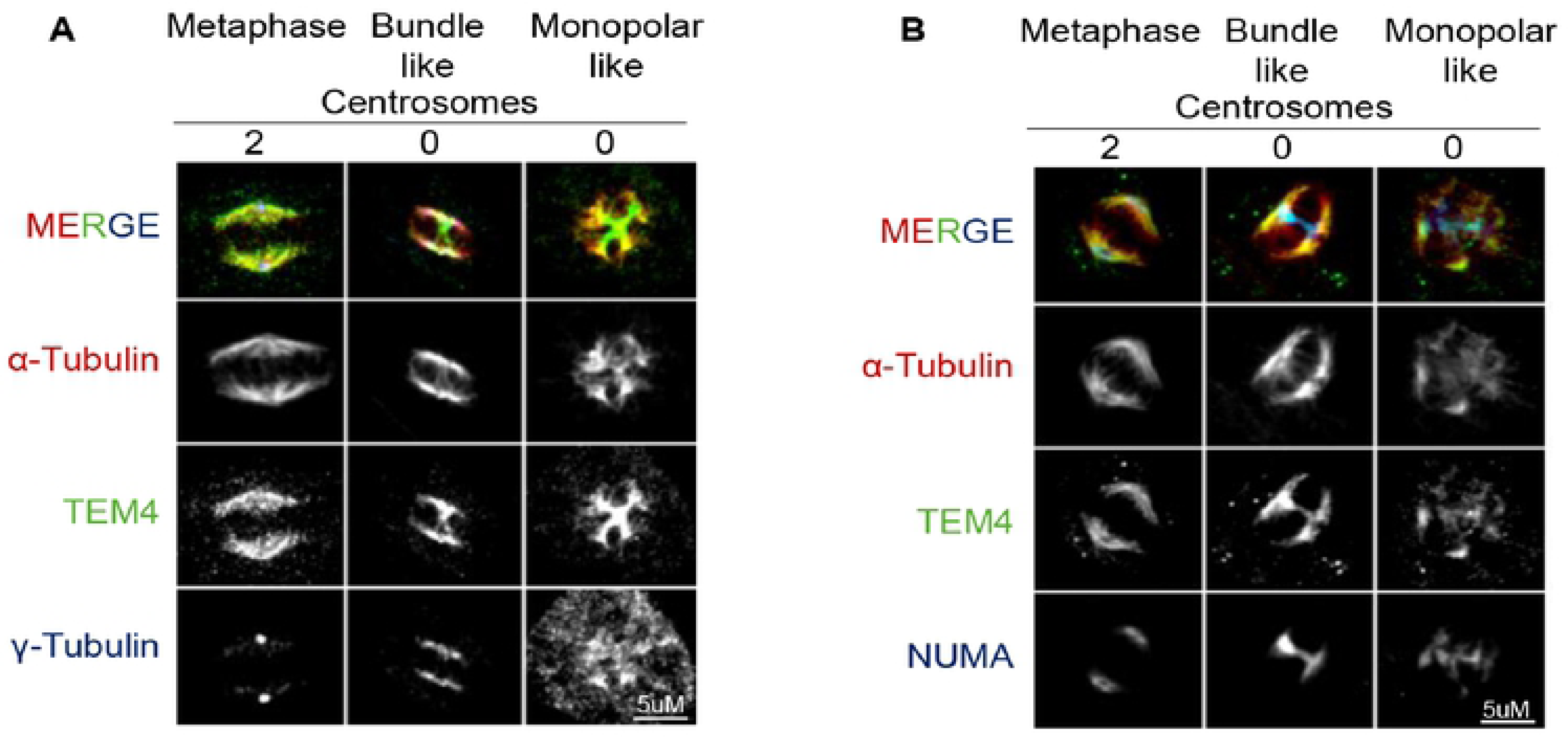
TEM4 and NuMA localize to similar microtubule structures in acentrosomal human cells. A: Immunofluorescent images of HeLa-T-REx cells treated with DMSO or 100 nM of centrinone for 72 hours. Cells were synchronized in mitosis after 10 hours of release from double thymidine block and fixed for 10 minutes with PTEMF buffer. Fixed cells were stained with anti-TEM4 (green), anti-α-tubulin for the spindle microtubules (red) anti-γ-tubulin (blue) to mark spindle poles. Scale bar = 5μm. B: Cells were treated as in A and then stained for NuMA (blue), TEM4 (green) and α- tubulin (red) in the presence or absence of centrinone as indicated. Scale bar = 5μm.

### TEM4 depletion results in reduction of TEM4 mRNA and protein in Western blotting but not the TEM4 signal by immunofluorescence

Our observations thus far suggest that TEM4 and NuMA may potentially colocalize and function together in spindle formation. In order to explore this idea in more detail, we generated inducible HeLa cell lines expressing two independently verified shRNAs targeting TEM4 (4,39,40). The efficiency of the depletion was initially evaluated by qPCR which demonstrated over 50% reduction of TEM4 mRNA in cells after 72 hours of shRNA induction compared to control cells (Fig. 5A). In parallel, and as a control, we show that TEM4 depletion did not significantly impact expression of NuMA mRNA levels (Fig. 5B). In agreement with the reduction in TEM4 mRNA levels, we observed a reduction of the TEM4 protein by Western blotting already after 48hours of induction, although the levels of the 180 kDa band remained unaffected (Fig. 5C). In stark contrast however, when we tested TEM4 depletion by immunofluorescence, we were unable to recapitulate these results and found no appreciable loss in the signal detected by the TEM4 antibody at the spindle, even after 72 hours of shRNA induction as shown in Fig. 5D and quantified in Fig.5E. Similarly, and as a control, we found no reduction of NuMA at the mitotic spindle. These data imply that the spindle signal detected by the commercial TEM4 antibody in immunofluorescence does not correspond to the TEM4 protein.

**Fig 5.**
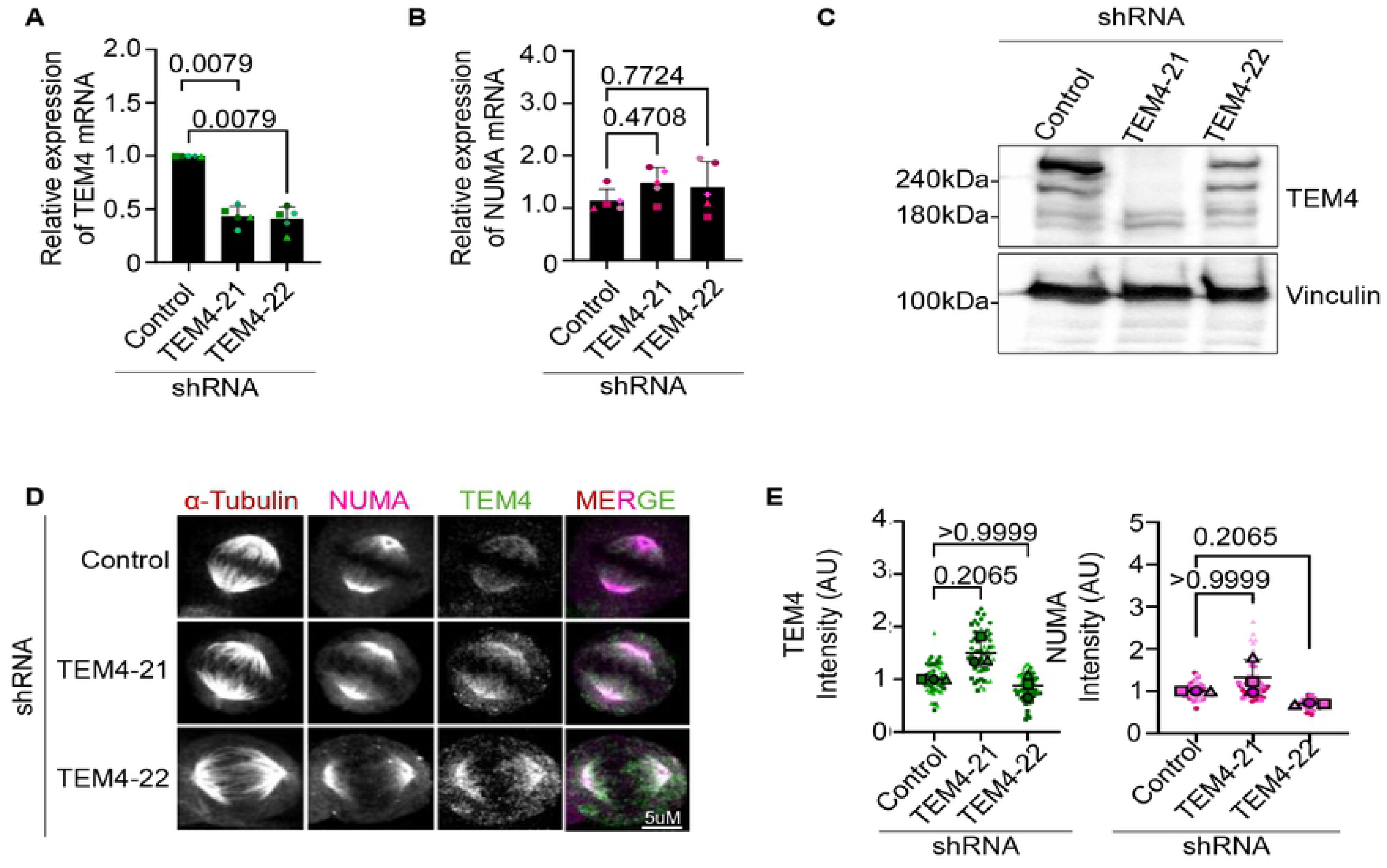
TEM4 depletion decreases TEM4 mRNA and protein levels but no effect on spindle staining by the TEM4 antibody. A: TEM4 gene expression levels on HeLa-T-REx cells transduced with lentiviruses for 72h expressing shRNA targeting TEM4. Levels of TEM4 mRNA were measured by RT-qPCR in 5 independent biological replicates. Gene expression of targeted mRNA was normalized to two reference genes, GAPDH and HPRT-1 B: Cells treated as in A, and NuMA mRNA levels were measured in Hela-T-REx cells depleted for TEM4. The graph shows the relative expression of NuMA mRNA from 5 independent biological replicates, normalized to two reference genes. C: HeLa-T-REx cells stably expressing two independent inducible shRNAs targeting TEM4 were generated. ShRNA expression was induced for 48 hours using 0.5 μg/ml doxycycline before total cell extraction with RIPA buffer followed and Western blotting was performed with the indicated antibodies D: Immunofluorescent images of HeLa-T-REx cells expressing shRNA to knockdown TEM4 after 72 hours of induction. Cells were synchronized in mitosis after 10 hours of release from double thymidine block and fixed for 10 minutes with PTEMF buffer. Fixed cells were stained with anti-TEM4 (green), anti-α-tubulin for the spindle microtubules (red) anti-NuMA (pink). Scale bar = 5μm. E: Cells were treated as in D and quantification of TEM4 and NuMA intensity on the mitotic spindle is shown in replicates from 3 independent experiments. A minimum of 20 cells was counted and the graphs show super plots of 3 biological replicates. One-way nonparamentrric Anova test was performed to compare the mean values between the tested groups.

### Depletion of NuMA results in loss of the immunofluorescence signal detected by the commercial TEM4 antibody

Considering the overlap between NuMA spindle localization and signals detected by the TEM4 antibody in immunofluorescence, we next asked if NuMA regulates TEM4 expression or localization. Efficient depletion of NuMA was achieved by siRNA using previously reported target sequences as shown by qPCR and Western blot analysis (Fig. 6A, B) (41) . NuMA depletion had no effect on TEM4 mRNA levels and we observed no decrease in the 220 kDa band corresponding to the predicted molecular mass of TEM4, although there was a marked reduction in the second 180 kDa band (Fig. 6C, D). Surprisingly, depletion of NuMA led to a drastic decrease of both NuMA and TEM4 antibody signal at the mitotic spindle as detected by immunofluorescence (Fig. 6E, F).

**Fig 6:**
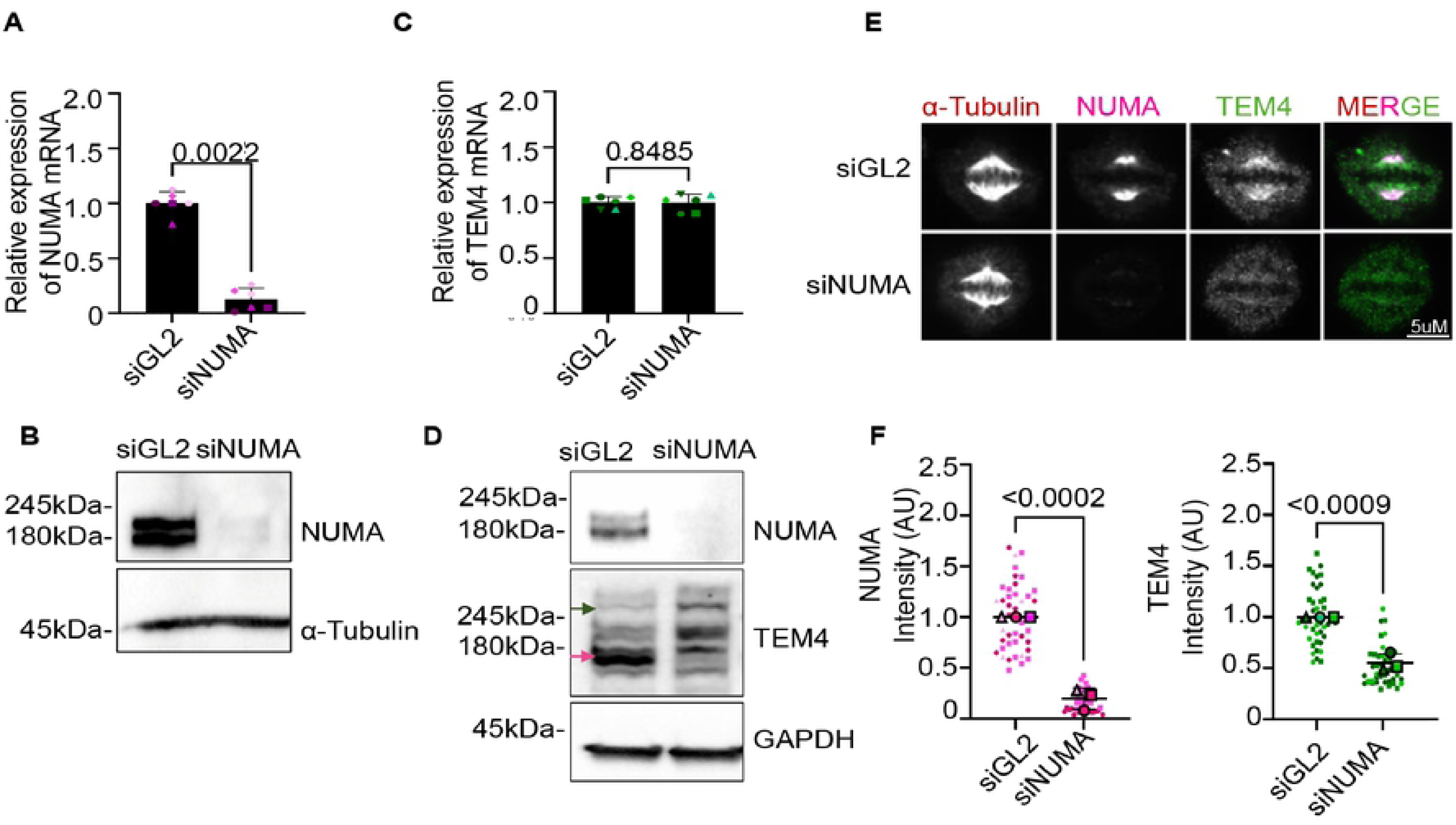
TEM4 microtubule staining is lost upon NuMA depletion without any alteration on the mRNA expression levels of TEM4. A: NuMA gene expression levels on HeLa-T-REx cells transfected for 72h with 120 nM of GL2 or NuMA siRNA. Levels of NuMA mRNA were measured by RT-qPCR in 5 independent biological replicates. Gene expression of targeted mRNA was normalized to two reference genes, GAPDH and HPRT-1 B: Cells were treated as in A. 72 hours post transfection total cells lysates were collected and total proteins were extracted with RIPA buffer. Lysates were subjected to Western Blot analysis and stained with α-tubulin as a loading control and NuMA C: TEM4 gene expression levels in HeLa-T-REx cells transfected with scrambled or NuMA siRNA for 72 hours. Levels of TEM4 mRNA were measured by RT-qPCR in 6 independent biological replicates. Gene expression of targeted mRNA was normalized to two reference genes, GAPDH and HPRT-1 D: Cells were treated as in C. 72 hours post transfection total cells lysates were collected and total proteins were extracted with RIPA buffer. Lysates were subjected to Western Blot analysis and immunoblotted with the indicated antibodies. E: Immunofluorescence images of HeLa-T-REx cells transfected with GL2 or NuMA siRNA for 72 hours as previously. Cells were synchronized in mitosis after 10 hours of release from double thymidine block and fixed for 10 minutes with PTEMF buffer. Fixed cells were stained with anti-TEM4 (green), anti-α-tubulin for the spindle microtubules (red) anti-NuMA (pink). Scale bar = 5μm. F: Quantification analysis of TEM4 and NuMA intensity at the mitotic spindle corresponding to 6E. A minimum of 15 cells per condition per experiment was counted and the graphs show superplots from 3 independent experiments. The P value is calculated from unpaired t-test

These data are in agreement with the idea that NuMA depletion does not decrease TEM4 mRNA or protein as detected by Western blotting, but drastically decreases the immunofluorescence signal detected at spindles by the TEM4 antibody.

### The TEM4 antibody recognizes both NuMA and TEM4

The loss of spindle staining detected by immunofluorescence and concomitant loss of the 180 kDa band detected by Western blotting with the TEM4 antibody upon NuMA depletion strongly suggests that this antibody cross-reacts with endogenous NuMA. To test this idea using an orthogonal approach, we performed immunoprecipitation and mass spectrometry to identify proteins associated with the TEM4 antibody. HeLa S3 cells were synchronized in mitosis after an overnight treatment with Taxol and mitotic cells were collected by shake-off. Samples from three independent experiments were lysed and an immunoprecipitation with either control IgG or TEM4 antibody was performed from equalized lysates. Considering all 3 replicates, the quantitative profile of identified proteins indicated 509 proteins common to both IgG and TEM4 samples, while only 23 proteins were uniquely identified in TEM4 immunoprecipitants (Fig. 7A). Analysis of peptide abundance in control and TEM4 antibody immunoprecipitants revealed a significant enrichment of normalized spectra corresponding to NuMA (with a mean of 85.00 ± 6.00 spectra per replicate) over TEM4 (mean of 37.67 ± 3.51 spectra per experiment, Fig 7B). In addition, the percentage of coverage for NuMA in TEM4 immunoprecipitates was 42± 1.52% while the percentage of TEM4 coverage was lower at 25± 1.73%. These results are in agreement with our previous observations and collectively support the idea that NuMA is strongly recognized by the TEM4 antibody by immunoprecipitation (Fig. 7B) and immunocytochemistry (Fig. 6E).

**Fig 7:**
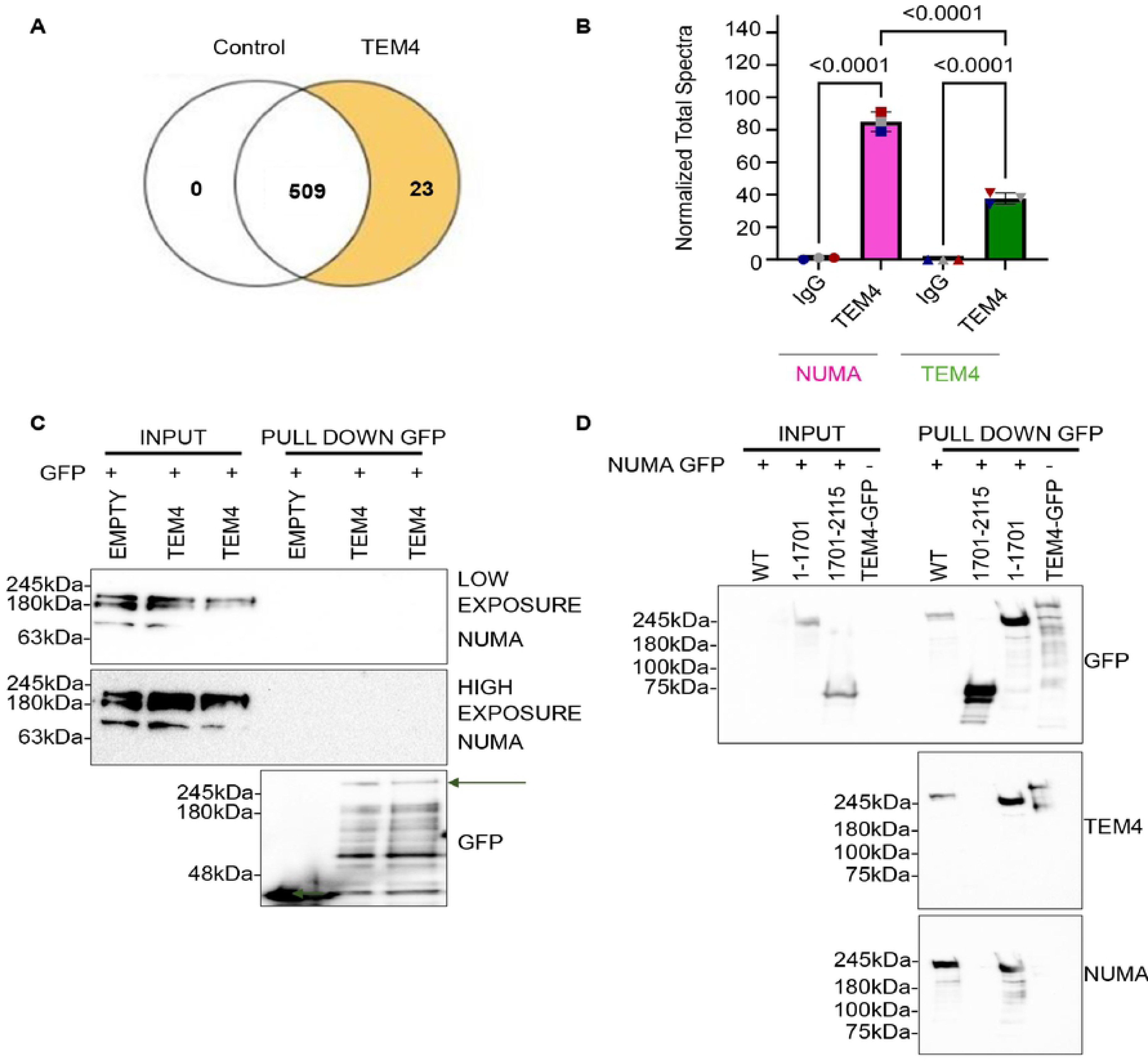
Validation of cross reactivity of TEM4 antibody with NuMA. A: Venn Diagram showing overlap in proteins identified in control samples, in TEM4 antibody immunoprecipitants or in both conditions. Control IgG and TEM4 and the identified proteins were annotated and normalized to the control samples. The 3 independent biological replicates identified 509 proteins detected in both IgG and TEM4 samples and 23 contained only in TEM4 samples and not at the control samples. B: Statistical analysis of normalized total spectra identified in control IgG samples and TEM4 samples. A t-test with Benjamini-Hochberg correction was performed, and the fold change was set by category with IgG as a reference category. C: Empty-GFP or GFP-TEM4 overexpression in HEK-293T cells, transiently transfected with FuGENE® HD Transfection Reagent according to manufactures’ protocol. Cells were treated with 15 nM Taxol 24 hours post transfection, to synchronize them in mitosis. Whole cell lysates were extracted with NP-40 buffer and immunoprecipitated (IP) with anti-GFP antibodies. Subsequently, samples were resolved by SDS-PAGE and immunoblotted with the indicated antibodies. D: Overexpressed GFP-NuMA constructs were immunoprecipitated with GFP antibody. HEK-293T cells were transiently transfected using FuGENE® HD Transfection Reagent according to manufactures’ protocol, with GFP-NuMA^FL^, GFP-NuMA^1-1701^ and GFP-NuMA^1701-2115^ alongside GFP-TEM4 as a positive control. Cells were synchronized in mitosis after 14 hours of treatment with 15nM of Taxol and total proteins were extracted with NP-40 buffer. GFP immunoprecipitates were immunoblotted with anti-GFP, anti- TEM4 and anti-NuMA antibodies.

A priori, co-immunoprecipitation of TEM4 and NuMA by the TEM4 antibody could occur if these proteins genuinely interact. To determine whether such an interaction exits without relying on the TEM4 antibody, we expressed either GFP or GFP-tagged TEM4 in HEK-293T cells and asked whether exogenous TEM4 can co-immunoprecipitate NuMA in mitotic extracts. Although NuMA was readily detected by Western blotting in total cell lysates, we failed to detect it in duplicate GFP-TEM4 immunoprecipitates, even after prolonged exposure, strongly suggesting that TEM4 and NuMA do not interact (Fig. 7C).

Our data thus far suggest that the commercial TEM4 antibody cross-reacts with NuMA by immunocytochemistry and in immunoprecipitates. Moreover, we speculate that the 180kDa band detected by this antibody in Western blotting and that is lost after NuMA depletion (Fig. 6D) corresponds to NuMA. To directly test this idea and determine whether the TEM4 antibody detects NuMA by Western blotting, we tested the capacity of the TEM4 antibody to detect different NuMA fragments corresponding to full-length NuMA (GFP- NuMA^FL^), NuMA residues 1-701 (GFP-NuMA^1-1701^) and NuMA residues 1701-2115 (GFP-NuMA^1701-2115^) alongside GFP-TEM4 as a control. GFP-immunoprecipitates were resolved by SDS-PAGE and immunoblotted with anti-GFP, -NuMA and -TEM4 antibodies. As shown in Fig. 7D, the TEM4 antibody detected not only TEM4 but also two of the three NuMA fragments, NuMA^FL^ and NuMA^1-1701^. Collectively, our data are in strong agreement with the idea that that the TEM4 antibody recognizes both NuMA and TEM4 by Western blotting and immunoprecipitation, whereas by immunofluorescence of mitotic cells, it reacts primarily with NuMA at the spindle.

## DISCUSSION

The need for better validation of the specificity and reproducibility of antibody reagents for biological research applications has been a much-discussed topic in recent literature (8–15). Here, we performed validation of a commonly used commercial TEM4 antibody (RRID: AB_1141641). Using a combination of multiple orthogonal approaches including immunoprecipiration and mass spectrometry, gene silencing, Western blotting, immunocytochemistry and antigen competition assays, we demonstrate that this antibody exhibits strong cross-reactivity, in particular by immunocytochemistry in mitotic cells, to the nuclear and spindle apparatus protein NuMA. We provide several lines of evidence in support of this argument. First, as previously reported, we found that plasmid-expressed TEM4 localized primarily to the actin cytoskeleton and actin-dependent structures such as cell junctions in interphase cells (3,4,17). Similarly, TEM4 localized predominantly to the actin cortex in mitosis when cells were fixed, and indirect immunofluorescence was performed under the same conditions (Fig. 1D). In contrast, staining mitotic cells with anti- TEM4 antibody revealed strong labelling of the mitotic spindle, but virtually no detectable signal at the cortex. Secondly, efficient depletion of TEM4 by two independent shRNAs (as detected by Western blotting and qPCR) failed to demonstrate a significant reduction in TEM4 antibody staining at the mitotic spindle, although this signal was essentially abolished in NuMA depleted cells (Fig. 6E). A parallel loss of the NuMA band was observed in NuMA-depleted protein lysates by Western blotting using this TEM4 antibody (Fig. 6D). Thirdly, mass spectrometry of immunoprecipitates generated using the TEM4 antibody identified both TEM4 and NuMA, with NuMA peptides significantly enriched over TEM4 peptides (Fig. 7B). Subsequent co-immunoprecipitation using GFP-TEM4 failed to demonstrate an interaction between the two proteins (Fig. 7C). Finally, using various exogenously expressed NuMA fragments, we find that that the TEM4 antibody can detect one or more epitopes in the first 1700 amino acids of NuMA by Western blotting (Fig. 7D).

TEM4 has recently emerged as an important player in cell cycle fidelity and having reliable tools will be critical to assessing its role effectively and accurately in this process. Interestingly, a recent study using this antibody on recombinant TEM4 demonstrated a direct interaction between TEM4 and MPS1, and provided evidence for TEM4 kinetochore localization by fluorescence cross correlation spectroscopy using tagged constructs, suggesting a direct role for TEM4 in mitosis (7). Similarly, using chromosome spreading techniques to remove spindle and cytoplasmic signals, staining with the TEM4 antibody suggested that a pool of this protein localizes to kinetochores; however, it will be important to validate this finding in cells depleted of TEM4. Although we did not observe kinetochore localization of overexpressed, tagged TEM4, we cannot exclude the presence of a small pool that is not easily detected by confocal microscopy.

TEM4 antibody RRID: AB_1141641 is a rabbit polyclonal antibody that was developed against a synthetic peptide targeting the middle region of TEM4 (residues 910 to 960). Alignment of this sequence with the first 1700 amino acids of NuMA identified a short uninterrupted stretch of moderate sequence similarity that may correspond the region of cross reactivity (Fig. S3), although it should be noted that antibody binding might be affected by protein conformation, and the epitope may be split in more than one peptide segment. Overall, our observations indicate that use of the anti-TEM4 antibody to detect TEM4, in particular using immunocytochemistry approaches, requires careful interpretation and controls.

## Materials and Methods

### Cell lines

HeLa S3, HeLa-T-REx Flp-IN, U20S, RPE-1, HEK-293T and HCT-116 cell lines were grown at 37°c with 5% CO2 in DMEM (Dulbecco’s Modified Eagle Medium, Hyclone) containing 10% fetal bovine serum or bovine growth serum (PAA) and supplemented with Penicillin-Streptomycin (100 µg/ml-1, Hyclone). For lentiviral induction, HeLa-T-REx cells were treated with 0.5 µg/ml of Doxycycline for 48 or 72 hours as specified.

### Drug treatments and transfections

Drug treatments were performed as follows unless otherwise indicated: Thymidine (Acros Organics, 2mM for 16 hours), MG132 (Calbiochem, 20μM for 2.5 hours), Nocodazole (Sigma, Cat# M1404-10MG, 100ng/mL for 12 hours), Centrinone (HY-18682, Cedarlane Labs, 100Nm for 72 hours), Taxol, (580555, Calbiochem, 15nM for 14 hours). Plasmids were transfected using FuGENE® HD Transfection Reagent according to manufacture’s protocol for 48 to 72 hours.

### siRNA knockdown

Endogenous protein depletion of NuMA was carried out with DsiRNAs (IDT), using JetPRIME® (Polyplus-transfection) for the HCT-116 and HeLa-T-REx cell. Cells were transfected at 50-60 % confluency and analyzed 72 hours post transfection.

The dsiRNAs target the following sequences in NuMA (41): dsiNuMA (used at 120nM) 5′ CAUGGCACUGAAGAGGGACAGCAAA −3′ and control dsiGL2: 5′- UCGAAGUAUUCCGCGUACG -3′

### ShRNA construction and generation of inducible cells lines

Inducible TEM4 knock down cell lines were generated as previously described (42, 43). Briefly, all shRNAs used in this study were cloned into EZ-Tet-pLKO PURO vector (Adgene 85966). The vector was prepared by digestion with NheI and EcoRI (NEB) for 4 hours at 37 °C. Cut vector was then dephosphorylated with 1μl of calf intestinal phosphatase (NEB) using the manufacturer’s protocol. shRNA oligos that were used have been previously validated and are the following (4,39,40): TRCN0000047521: 5′- CCGGGTATCTGAATAACCAGGTGTTCTCGAGAACACCTGGTTATTCAGATACTTTTTG-3′ and TRCN0000047522 : 5′- CCGGGAGGTTATTCAGAGCATAGTTCTCGAGAACTATGCTCTGAATAACCTCTTTTTG-3′. Oligos were annealed in buffer containing 10mM TRIS pH 7.4 and, 1mM EDTA and 1M NaCl in the following PCR program (95 °C 10 mins, 95 °C to 85 °C -2 °C/s, 85 °C 1 min, 85 °C to 75 °C -0.3 °C/s, 75 °C 1 min, 75 °C to 65 °C -0.3 °C/s, 65 °C 1 min, 65 °C to 55 °C -0.3 °C/s, 55 °C 1 min, 55 °C to 45 °C -0.3 °C/s, 45 °C 1 min, 45 °C to 35 °C -0.3 °C/s, 35 °C 1 min, 35 °C to 25 °C -0.3 °C/s, 25 °C 1 min, 4 °C hold). Ligation of prepared vector and oligos was performed with T4 Ligase (NEB) overnight at 16 °C. PCR screening followed to confirm the right clones. Constructs were used to make lentivirus in HEK-293T cells using the ViraPower system (K497500, Invitrogen). Plates were coated with 2 μg/mL PolyD lysine in PBS (phosphate-buffered saline) for 1 hour at 37 °C before cells were seeded. 24 hours post seeding; media was replaced by antibiotic-free media with heat-inactivated serum and the transfection was performed with Lipofectamine2000 (ThermoFisher). Media was changed to the target cell media (without antibiotics) after 24 hours and cells were then returned to 37 °C for 48 hours to produce viral particles. Viral media was collected and centrifuged for 10 minutes at 1500 RPM to pellet cell debris. Next, the viral media was filtered by syringe through a 0.45 μM, low protein binding filter (28145–505, VWR). Cells were typically infected by first adding half the volume with normal growth media (no antibiotics, heat inactivated serum) and half volume with the filtered viral media plus polybrene to a 5 μg/mL final concentration to improve infection rate. Infected cells were incubated 48-72 hours and then given fresh growth media for an additional 24-48 hours before beginning selection. After lentivirus preparation and infection, single cell clones were expanded and screened via immunoblot and genomic sequencing.

### Microtubule regrowth assay

Microtubule regrowth assay was performed as previously described (44). Briefly cells were plated two days before fixation. Next, cells were treated with 3.3 μM of Nocodazole for three hours. Media containing Nocodazole was washed out 4 times with warm PBS, followed by a wash in warm media and incubation of cells in worm media at indicated time points to allow the regrowth of microtubules. Cold methanol was added to the cells for 10 minutes at – 20 °C for simultaneous fixation and permeabilization.

### Cold treatment assay

For cold-treatment assays, cells were washed twice with PBS, incubated on ice for 20 minutes in the presence of cold DMEM medium and washed once with cold PBS before fixation. Fixation was performed with cold PTEMF buffer (0.2% Triton X-100, 20mM PIPES pH 6.8, 1mM MgCl2, 10mM EGTA and 4% formaldehyde) for 10 minutes.

### Mass spectrometry

For Mass spectrometry experiments HeLa-T-REx cells were grown in 150 mm plates and treated with 15 nM of Taxol for 16 hours. Mitotic cells were collected by mitotic shake off and lysed in NP40 buffer (150 mM NaCl, 1.0% NP-40, 50 mM Tris-cl, pH8%). BCA quantification was performed, and 10 mg of protein was used for the immunoprecipitation (IP) with 10 μg of TEM4 antibody or 10 μg of IgG. The IP was performed at 4°C for 2 hours followed by 50 μl of Protein G beads, washed 3 times in NP40 buffer. Lysates and beads were incubated at 4°C for 1 hour, washed twice with NP-40 buffer and 5 times with 20 mM of Ammonium Bicarbonate. Peptide digestion was performed without alkylation. Briefly, equal bead volume of diluted trypsin (1ng/ml) was added to the beads and incubated at 37 °C for 5 hours. Trypsin digestion was stopped by addition of 1% formic acid solution. Supernatant was harvested, and beads were resuspended in 60% acetonitrile with 0,1 formic acid at room temperature for 5 minutes. Samples were dried in a speed vac and resuspended in 20 μl of 0.1 trifluoroacetic acid (TFA) to desalt on ZipTip (Thermo Scientific Pierce, PI87782) before mass spectrometry. Mass spectrometry analysis was performed by the Proteomics Platform of the CHU de Québec Research Center (Quebec, Qc, Canada). Samples were analyzed by nano LC-MS/MS using a Dionex UltiMate 3000 nanoRSLC chromatography system (Thermo Fisher Scientific) connected to an Orbitrap Fusion mass spectrometer (Thermo Fisher Scientific, San Jose, CA, USA). Peptides were trapped at 20 μl/min in loading solvent (2% acetonitrile, 0.05% TFA) on a 5mm x 300 μm C18 pepmap cartridge pre-column (Thermo Fisher Scientific / Dionex Softron GmbH, Germering, Germany) during 5 minutes. Then, the pre-column was switched online with a Pepmap Acclaim column (ThermoFisher) 50 cm x 75µm internal diameter separation column and the peptides were eluted with a linear gradient from 5-40% solvent B (A: 0,1% formic acid, B: 80% acetonitrile, 0.1% formic acid) in 60 minutes, at 300 nL/min for a total run time of 90 minutes. Mass spectra were acquired using a data dependent acquisition mode using Thermo XCalibur software version 4.3.73.11. Full scan mass spectra (350 to 1800m/z) were acquired in the orbitrap using an AGC target of 4e5, a maximum injection time of 50 ms and a resolution of 120 000. Internal calibration using lock mass on the m/z 445.12003 siloxane ion was used. Each MS scan was followed by MS/MS fragmentation of the most intense ions for a total cycle time of 3 seconds (top speed mode). The selected ions were isolated using the quadrupole analyzer in a window of 1.6 m/z and fragmented by Higher energy Collision-induced Dissociation (HCD) with 35% of collision energy. The resulting fragments were detected by the linear ion trap in rapid scan rate with an AGC target of 1e4 and a maximum injection time of 50ms. Dynamic exclusion of previously fragmented peptides was set for a period of 20 sec and a tolerance of 10 ppm.

### Database searching

MGF peak list files were created using Proteome Discoverer 2.5.0software (Thermo). MGF sample files were then analyzed using Mascot (Matrix Science, London, UK; version 2.8.0). Mascot was set up to search a contaminant database and REF_HomoSapiens_20200510_UP000005640_20200924 database (unknown version, 97134 entries) assuming the digestion enzyme trypsin. Mascot was searched with a fragment ion mass tolerance of 0.60 Da and a parent ion tolerance of 10.0 PPM. Carbamidomethyl of cysteine was specified in Mascot as a fixed modification. Deamidation of asparagine and glutamine and oxidation of methionine were specified in Mascot as variable modifications. 2 missed cleavages were allowed.

Data analysis was performed using Scaffold (version Scaffold_5.1.0, Proteome Software Inc., Portland, OR) to validate MS/MS based peptide and protein identifications. Peptide identifications were accepted if they could be established at greater than 95% probability using the Scaffold delta-mass correction. Protein identifications were accepted if they contained less than 1% false discovery rate (FDR) and least two peptides. All results from the 3 different biological replicates, were organized in two different categories, control IgG and TEM4, and quantitative analysis on normalized total spectra was performed. Statistical significance was determined by t-test and Benjamini-Hocheberg correction.

### Protein extraction

Lysis of cell pellets for immunoblotting and immunoprecipitation was done with RIPA lysis buffer (150mM Tris-HCL pH 7.5, 150mM NaCl, 10mM NaF, 1% NP-40, 0.1% Na-deoxycholate and a protease and phosphatase inhibitor cocktail that included 20mM B-glycerophosphate, 0.1mM sodium vanadate, 10mM sodium pyrophosphate, 1mg.ml-1, leupeptin, 1mg.ml-1, aprotinin and 1mM AEBSF) or 8M Urea lysis buffer (8M urea, 50 mM HEPES, 5% glycerol, 1.5mM MgCl2 and the same protease and phosphatase inhibitor cocktail used in the RIPA buffer). Cells were lysed on an orbital shaker at 4°C for minimum 30 minutes and lysates were centrifuged at maximum speed for 20 minutes at 4°C. The supernatant was collected and quantified using the BCA assay (Thermo Scientific) for protein concentration.

### Western-Blot

Between 40 and 80 µg of proteins were loaded onto 8-12% SDS-PAGE gels. Proteins were migrated and then transferred onto a PVDF transfer membrane (0.45 micron, Immobilon-P) using overnight wet transfer. Membranes were blocked in 5% of milk diluted in PBS and containing 0.05% of Tween 20 for 30 minutes, and then, incubated overnight at 4°C with specific primary antibody in 5% of milk in PBS. Membranes were washed with PBS containing 0.05% of Tween-20 and incubated with the appropriate secondary antibody conjugated to horseradish peroxidase for 1 hour at room temperature. After three additional washes, antibody binding was detected with either the Clarity or Clarity Max Western ECL substrate (Bio-Rad) and ChemiDoc MP Imaging System (Bio-Rad).

### RNA extraction and purification

Total RNA was extracted from HeLa-T-REx cells after incubation in RLT lysis buffer (Qiagen) containing 10% of β-mercaptoethanol (Sigma, M3148) for lysing cells prior to RNA isolation, and purified with the RNeasy Mini Kit (Qiagen) according to the manufacturer’s protocol. In order to avoid potential genomic contamination, samples were incubated with RNase-free DNase (Qiagen). The quantity and the quality of purified total RNA was assessed on a NanoDrop 1000 microvolume spectrophotometer (Thermo Scientific). Samples were stored at −80°C until use.

### Reverse transcription and quantitative real-time PCR (qRT-PCR)

iScript Advanced cDNA Synthesis Kit for PCR (BioRad) was used for Reverse Transcription (RT) of one µg for each sample, following the manufacturer’s protocol. Real-time quantitative PCR was carried out on cDNA samples from HeLa-T-REx cells using SsoAdvanced Universal SYBR Green Supermix (BioRad). The primers sequence was the following: hGAPDH: Forward 5’-GAAAGCCTGCCGGTGACTAA-3’, Reverse 5’- GCCCAATACGACCAAATCAGAGA-3’, hHPRT1: Forward 5’- TGCTGAGGATTTGGAAAGGGT-3’, Reverse 5’-AGCAAGACGTTCAGTCCTGT-3’, TEM4: Forward 5’-TACATGCTGAACCTGCACTCC-3’, Reverse 5’GTGCTTCCGCATGTCCACC-3’, NUMA: Forward 5’-AGCCAGACTGCTGGAGATCA-3’, Reverse 5’- CCCTCTAGCACCTTCTGTGC-3’. 0.5 μM of forward and reversed primer were combined with 2 μL cDNA samples, for the qRT-PCR. Two negative controls were included: an RT-negative control and a no-template control. Samples were incubated at 95°C for 5 minutes followed by 40 cycles of three amplification steps: 95°C for 15 seconds, the optimal primer-specific temperature (between 50 and 60°C) for 15 seconds and 72°C for 15 seconds. Each qRT-PCR reaction was performed as two technical replicates for each biological sample, and then, normalized to two reference genes: GAPDH and HPRT. Results were analyzed using the Pfaffl method to calculate fold inductions and express the results as relative quantification values based on cycle threshold (Ct) comparisons between different samples (45). Experiments were performed on 5 to 6 biological replicates to validate the expression levels of TEM4 and NuMA.

### Immunofluorescence and antibodies

Cells were grown on coverslips coated with 25µg/ml poly-ethylenimine (PEI) and 150mM NaCl solution and treated with indicated drugs or dsiRNA after adhesion. Cells were fixed in PTEMF buffer for 10 minutes at room temperature. After fixation, coverslips were blocked with 3% bovine serum albumin (BSA) in PBS-Tween 0.2% for at least 30 minutes. Coverslips were incubated with primary and secondary antibodies for 2 hours and 1 hour respectively at room temperature. Antibodies were used at 1 µg/ml, unless otherwise indicated, as follows: -anti-TEM4 (ab67278, Abcam), anti-α-tubulin (DM1A, Santa Cruz), CREST anti-centromere serum (HCT-0100, Immunovision), anti-NuMA (GT3611, Thermo Scientific), anti-GFP (1:500, Millpore), anti-γ-tubulin ([TU-30] ab27074, Abcam), anti-GAPDH ((1A10) (NBP1-47339, Novus Biologicals)), anti-phalloidin–Atto 565 (9402, Sigma Aldrich). To stain chromosomes, HOESCHT 33342 (Sigma) was used. Alexa Fluor-AffiniPure series secondary antibodies (Thermo) or (Jackson ImmunoResearch) were used for immunofluorescence (1:1000) and horseradish peroxidase-coupled secondary antibodies (Jackson ImmunoResearch) were used for western blotting (1:10000).The peptide used in in the competition assay was purchased from Novus Biologicals (Cat. number: NBP1-77314PEP).

### Microscopy

All images were acquired by confocal microscopy on an inverted Olympus IX80 microscope equipped with a WaveFX-Borealin-SC Yokagawa spinning disc (Quorum Technologies) and an Orca Flash4.0 camera (Hamamatsu). Images were acquired by Metamorph software (Molecular Devices). Optical sections were acquired with identical exposure times for each channel within an experiment and then projected into a single picture using ImageJ (rsb.info.nih.gov). Image J was used for image processing and all images shown in the same figure have been identically scaled.

### Quantification and statistical analysis

Unless otherwise stated, all experiments and statistical analysis were performed on triplicate. Image quantification was realized using Image J. For measurement of signal intensities at the mitotic spindle, the α-tubulin signal was used to generate a binary mask. Integrated signal intensity was measured in all relevant channels and intensities indicated are values relative to a tubulin or arbitrary units. A minimum of 20 cells was quantified per condition for all experiments. Statistical analysis and graphic plotting were performed in GraphPad Prism Software V6.01. Non-parametric t test (Mann-Whitney test) was used in experiments comparing only two conditions, while ANOVA (Kruskal-Wallis test) was used for experiments with more than two conditions compared. Statistical significance was considered with a P-value smaller than 0.005

## ACKNOWLEDGEMENTS

We thank Dr. Jan Ellenberg for candid discussions. This work was supported by Canadian Institutes of Health Research grants to S.E. (project grants 376557 and 388292). S.E. holds an FRQS (Fonds de Recherche de Santé Québec) Senior researcher salary award. D.K.P has been supported by training awards from Desjardins, PROTEO, and by a CRCHU de Québec training award.

---

**Fig.S1.**
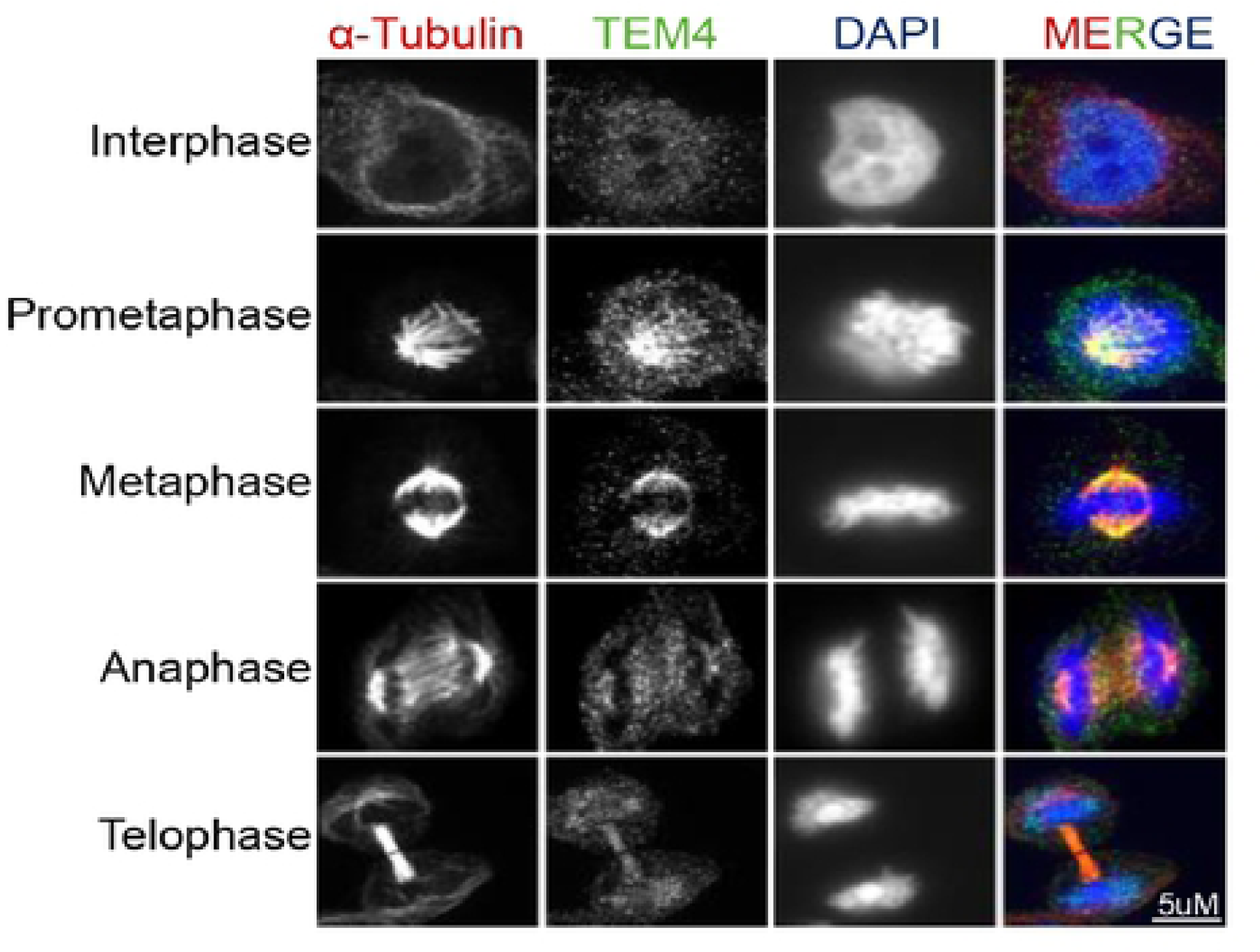
The TEM4 antibody decorates the mitotic spindle in HeLa-T-Rex. Representative immunofluorescence image of metaphase spindle HeLa-T-REx cell. Cells were synchronized in mitosis after 10 hours of release from double thymidine block and fixed for 10 minutes with PTEMF buffer. Fixed cells were stained with anti- TEM4 (green), anti-α-tubulin for the spindle microtubules (red) and HOECHST to stain the DNA (blue). Scale bar = 5μm.

**Fig.S2.**
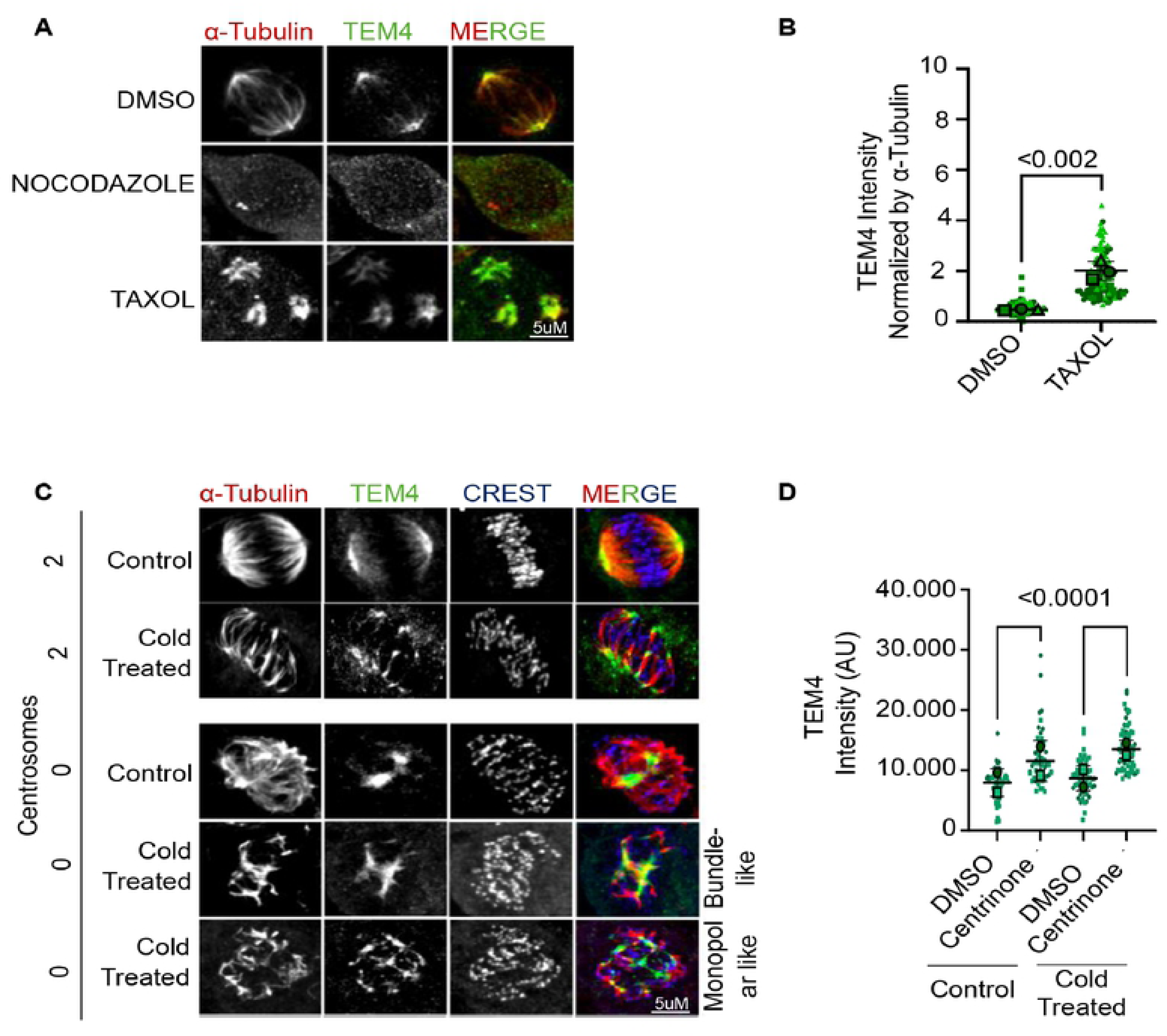
The TEM4 antibody staining is altered after disturbing microtubule stability. A: Immunofluorescence image of HCT-116 cells treated with 3 μM of Nocodazole or 15 nM of Taxol for 14 hours. Cells were fixed for 10 minutes with PTEMF buffer and stained with anti-TEM4 (green), anti-α-tubulin for the spindle microtubules (red). Scale bar = 5μm. B: Superplot of the quantification of TEM4 intensity at the mitotic spindle in 3 independent experiments corresponding to A. A minimum of 20 cells was counted per condition in each experiment. The P value is calculated using the unpaired t-test. C: Immunofluorescent images of HeLa-T-REx cells treated with DMSO or 100 nM of centrinone for 72 hours. Cells were synchronized in mitosis after 10 hours of release from double thymidine block and 20 minutes before fixation cells were incubated on ice to depolymerize microtubules. Fixation was performed by 10 minutes incubation with PTEMF buffer. Fixed cells were stained with anti-TEM4 (green), anti-α-tubulin for the spindle microtubules (red) and CREST serum for the kinetochores (blue). Scale bar = 5μm. D: Superplot graphs showing quantification of TEM4 intensity at the mitotic spindle corresponding to C. A minimum of 20 cells was counted and two biological replicates were analyzed.

**Fig.S3.**
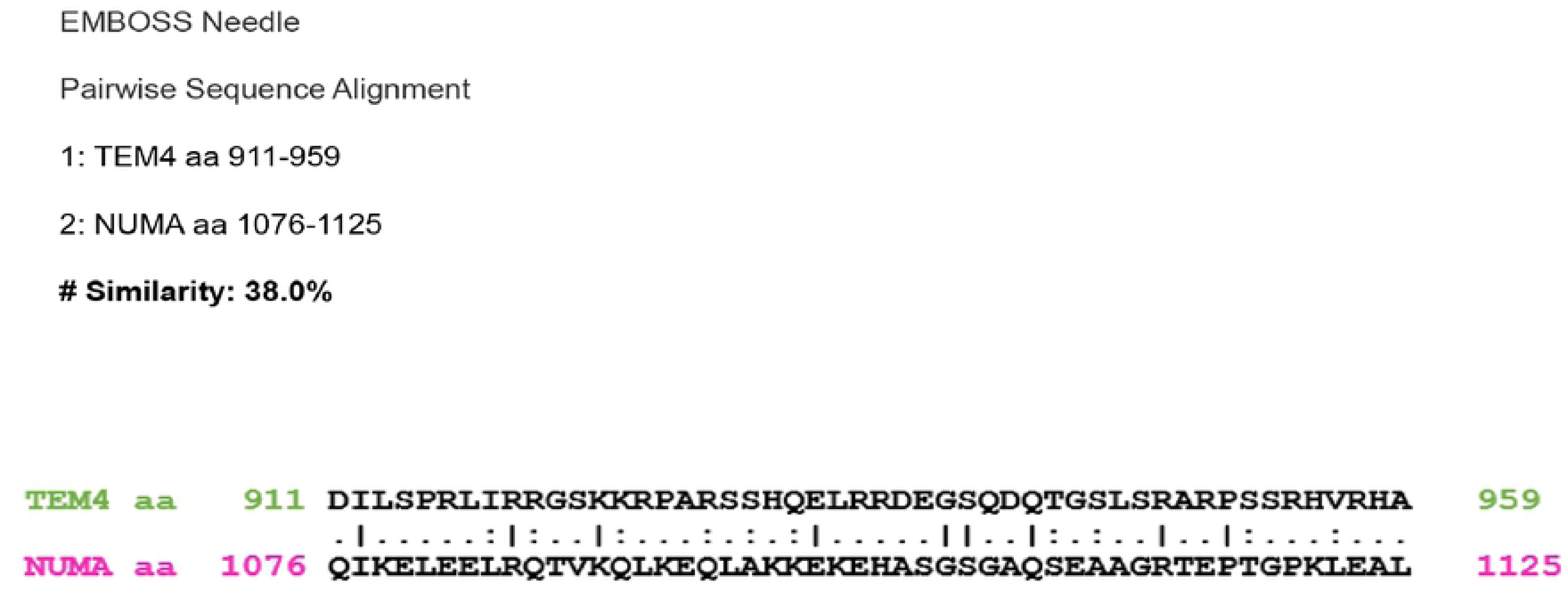
Alignment of the TEM4 antibody antigen with a potentially overlapping region of NuMA. Alignment of TEM4 sequence (residues 911 to 959), (Q96PE2|ARHGH_HUMAN Rho guanine nucleotide exchange factor) and NuMA sequence (residues 1 to 1125), (Q14980|NuMA1_HUMAN Nuclear mitotic apparatus protein 1) using EMBOSS Needle. A pairwise sequence alignment was performed, and the percentage of similarity for those sequences was 38%

## REFERENCES

1. Hall A. Rho family GTPases. Biochemical Society Transactions. 2012 Nov 21;40(6):1378–82.

2. Jaffe AB, Hall A. Rho GTPases: biochemistry and biology. Annu Rev Cell Dev Biol. 2005;21:247–69.

3. Mitin N, Rossman KL, Currin R, Anne S, Marshall TW, Bear JE, et al. The RhoGEF TEM4 Regulates Endothelial Cell Migration by Suppressing Actomyosin Contractility. PLoS One. 2013 Jun 18;8(6):e66260.

4. Ngok SP, Geyer R, Kourtidis A, Mitin N, Feathers R, Der C, et al. TEM4 is a junctional Rho GEF required for cell–cell adhesion, monolayer integrity and barrier function. J Cell Sci. 2013 Aug 1;126(15):3271–7.

5. García-Jiménez I, Cervantes-Villagrana RD, del-Río-Robles JE, Castillo-Kauil A, Beltrán-Navarro YM, García-Román J, et al. Gβγ mediates activation of Rho guanine nucleotide exchange factor ARHGEF17 that promotes metastatic lung cancer progression. Journal of Biological Chemistry [Internet]. 2022 Jan 1 [cited 2022 Jan 18];298(1). Available from: https://www-jbc-org.acces.bibl.ulaval.ca/article/S0021-9258(21)01249-7/abstract

6. Neumann B, Walter T, Hériché JK, Bulkescher J, Erfle H, Conrad C, et al. Phenotypic profiling of the human genome by time-lapse microscopy reveals cell division genes. Nature. 2010 Apr;464(7289):721–7.

7. Isokane M, Walter T, Mahen R, Nijmeijer B, Hériché JK, Miura K, et al. ARHGEF17 is an essential spindle assembly checkpoint factor that targets Mps1 to kinetochores. J Cell Biol. 2016 Mar 14;212(6):647–59.

8. Baker M. When antibodies mislead: the quest for validation. Nature. 2020 Sep 7;585(7824):313–4.

9. Baker M. Reproducibility crisis: Blame it on the antibodies. Nature. 2015 May 1;521(7552):274–6.

10. Cutts EE, Taylor GC, Pardo M, Yu L, Wills JC, Choudhary JS, et al. A commercial antibody to the human condensin II subunit NCAPH2 cross-reacts with a SWI/SNF complex component. Wellcome Open Res. 2021 Jan 13;6:3.

11. Edfors F, Hober A, Linderbäck K, Maddalo G, Azimi A, Sivertsson Å, et al. Enhanced validation of antibodies for research applications. Nat Commun. 2018 Oct 8;9(1):4130.

12. Forsström B, Axnäs BB, Stengele KP, Bühler J, Albert TJ, Richmond TA, et al. Proteome-wide Epitope Mapping of Antibodies Using Ultra-dense Peptide Arrays. Mol Cell Proteomics. 2014 Jun;13(6):1585–97.

13. Gokula M, Holmes HM. Tools to Reduce Polypharmacy. Clinics in Geriatric Medicine. 2012 May 1;28(2):323–41.

14. Lukinavičius G, Lavogina D, Gönczy P, Johnsson K. Commercial Cdk1 antibodies recognize the centrosomal protein Cep152. Biotechniques. 2013 Sep;55(3):111–4.

15. Petry FR, Pelletier J, Bretteville A, Morin F, Calon F, Hébert SS, et al. Specificity of Anti-Tau Antibodies when Analyzing Mice Models of Alzheimer’s Disease: Problems and Solutions. PLOS ONE. 2014 May 2;9(5):e94251.

16. Uhlen M, Bandrowski A, Carr S, Edwards A, Ellenberg J, Lundberg E, et al. A proposal for validation of antibodies. Nat Methods. 2016 Oct;13(10):823–7.

17. Mitin N, Rossman KL, Der CJ. Identification of a Novel Actin-Binding Domain within the Rho Guanine Nucleotide Exchange Factor TEM4. PLoS One. 2012 Jul 24;7(7):e41876.

18. Asiedu M, Wu D, Matsumura F, Wei Q. Phosphorylation of MyoGEF on Thr-574 by Plk1 Promotes MyoGEF Localization to the Central Spindle *. Journal of Biological Chemistry. 2008 Oct 17;283(42):28392–400.

19. Birkenfeld J, Nalbant P, Bohl BP, Pertz O, Hahn KM, Bokoch GM. GEF-H1 Modulates Localized RhoA Activation during Cytokinesis under the Control of Mitotic Kinases. Developmental Cell. 2007 May;12(5):699–712.

20. Cannet A, Schmidt S, Delaval B, Debant A. Identification of a mitotic Rac-GEF, Trio, that counteracts MgcRacGAP function during cytokinesis. Mol Biol Cell. 2014 Dec 15;25(25):4063–71.

21. Rodriguez-Fraticelli AE, Vergarajauregui S, Eastburn DJ, Datta A, Alonso MA, Mostov K, et al. The Cdc42 GEF Intersectin 2 controls mitotic spindle orientation to form the lumen during epithelial morphogenesis. J Cell Biol. 2010 May 17;189(4):725–38.

22. De Brabander M, Geuens G, Nuydens R, Willebrords R, De Mey J. Taxol induces the assembly of free microtubules in living cells and blocks the organizing capacity of the centrosomes and kinetochores. Proc Natl Acad Sci U S A. 1981 Sep;78(9):5608–12.

23. Schiff PB, Fant J, Horwitz SB. Promotion of microtubule assembly in vitro by taxol. Nature. 1979 Feb;277(5698):665–7.

24. Job D, Fischer EH, Margolis RL. Rapid disassembly of cold-stable microtubules by calmodulin. Proc Natl Acad Sci U S A. 1981 Aug;78(8):4679–82.

25. Stearns T, Kirschner M. In vitro reconstitution of centrosome assembly and function: the central role of gamma-tubulin. Cell. 1994 Feb 25;76(4):623–37.

26. Knop M, Pereira G, Geissler S, Grein K, Schiebel E. The spindle pole body component Spc97p interacts with the gamma-tubulin of Saccharomyces cerevisiae and functions in microtubule organization and spindle pole body duplication. EMBO J. 1997 Apr 1;16(7):1550–64.

27. Zheng Y, Wong ML, Alberts B, Mitchison T. Nucleation of microtubule assembly by a γ-tubulin-containing ring complex. Nature. 1995 Dec;378(6557):578–83.

28. Kollman JM, Merdes A, Mourey L, Agard DA. Nature Reviews Molecular Cell Biology Microtubule nucleation by γ-tubulin complexes. Nat Rev Mol Cell Biol. 2011 Oct 12;12(11):709–21.

29. Sulimenko V, Hájková Z, Klebanovych A, Dráber P. Regulation of microtubule nucleation mediated by γ-tubulin complexes. Protoplasma. 2017 May 1;254(3):1187–99.

30. Farache D, Emorine L, Haren L, Merdes A. Assembly and regulation of γ-tubulin complexes. Open Biol. 2018 Mar 7;8(3):170266.

31. Wieczorek M, Urnavicius L, Ti SC, Molloy KR, Chait BT, Kapoor TM. Asymmetric molecular architecture of the human γ-tubulin ring complex. Cell. 2020 Jan 9;180(1):165–175.e16.

32. Khodjakov A, Cole RW, Oakley BR, Rieder CL. Centrosome-independent mitotic spindle formation in vertebrates. Current Biology. 2000 Jan 15;10(2):59–67.

33. Maiato H, Rieder CL, Khodjakov A. Kinetochore-driven formation of kinetochore fibers contributes to spindle assembly during animal mitosis. J Cell Biol. 2004 Dec 6;167(5):831–40.

34. Meunier S, Vernos I. Acentrosomal Microtubule Assembly in Mitosis: The Where, When, and How. Trends in Cell Biology. 2016 Feb 1;26(2):80–7.

35. Roostalu J, Surrey T. Microtubule nucleation: beyond the template. Nat Rev Mol Cell Biol. 2017 Nov;18(11):702–10.

36. Sanchez AD, Feldman JL. Microtubule-organizing centers: from the centrosome to non-centrosomal sites. Curr Opin Cell Biol. 2017 Feb;44:93–101.

37. Wong YL, Anzola JV, Davis RL, Yoon M, Motamedi A, Kroll A, et al. Reversible centriole depletion with an inhibitor of Polo-like kinase 4. Science. 2015 Jun 5;348(6239):1155–60.

38. Chinen T, Yamamoto S, Takeda Y, Watanabe K, Kuroki K, Hashimoto K, et al. NuMA assemblies organize microtubule asters to establish spindle bipolarity in acentrosomal human cells. EMBO J. 2020 Jan 15;39(2):e102378.

39. Memon D, Gill MB, Papachristou EK, Ochoa D, D’Santos CS, Miller ML, et al. Copy number aberrations drive kinase rewiring, leading to genetic vulnerabilities in cancer. Cell Rep. 2021 May 18;35(7):109155.

40. Weber P, Baltus D, Jatho A, Drews O, Zelarayan LC, Wieland T, et al. RhoGEF17—An Essential Regulator of Endothelial Cell Death and Growth. Cells. 2021 Apr;10(4):741.

41. Iwakiri Y, Kamakura S, Hayase J, Sumimoto H. Interaction of NuMA protein with the kinesin Eg5: its possible role in bipolar spindle assembly and chromosome alignment. Biochemical Journal. 2013 Apr 15;451(2):195–204.

42. Frank SB, Schulz VV, Miranti CK. A streamlined method for the design and cloning of shRNAs into an optimized Dox-inducible lentiviral vector. BMC Biotechnol. 2017 Feb 28;17:24.

43. Moffat J, Grueneberg DA, Yang X, Kim SY, Kloepfer AM, Hinkle G, et al. A Lentiviral RNAi Library for Human and Mouse Genes Applied to an Arrayed Viral High-Content Screen. Cell. 2006 Mar 24;124(6):1283–98.

44. Tulu US, Fagerstrom C, Ferenz NP, Wadsworth P. Molecular requirements for kinetochore-associated microtubule formation in mammalian cells. Curr Biol. 2006 Mar 7;16(5):536–41.

45. Pfaffl MW. A new mathematical model for relative quantification in real-time RT– PCR. Nucleic Acids Res. 2001 May 1;29(9):e45.

